# Association between erythrocyte dynamics and vessel remodelling in developmental vascular networks

**DOI:** 10.1101/2020.05.21.106914

**Authors:** Qi Zhou, Tijana Perovic, Ines Fechner, Lowell T. Edgar, Peter R. Hoskins, Holger Gerhardt, Timm Krüger, Miguel O. Bernabeu

## Abstract

Sprouting angiogenesis is an essential vascularisation mechanism consisting of sprouting and remodelling. The remodelling phase is driven by rearrangements of endothelial cells (ECs) within the post-sprouting vascular plexus. Prior work has uncovered how ECs polarise and migrate in response to flow-induced wall shear stress (WSS). However, the question of how the presence of erythrocytes (well-known as RBCs) and their haemodynamics impact affects vascular remodelling remains unanswered. Here, we devise a computational framework to model cellular blood flow in developmental mouse retina. We demonstrate a previously unreported highly heterogeneous distribution of RBCs in primitive vasculature. Furthermore, we report a strong association between vessel regression and RBC depletion, and identify plasma skimming as the driving mechanism. Live imaging in a developmental zebrafish model confirms this association. Taken together, our results indicate that RBC dynamics are fundamental to establishing the regional WSS differences driving vascular remodelling via their ability to modulate effective viscosity.

**Summary:** Recent studies demonstrate that during sprouting angiogenesis, blood flow provides crucial hydrodynamic cues (*e.g.* wall shear stress) for the remodelling of primitive plexuses towards a functional network. Notwithstanding, the role of RBCs in this process remains poorly understood. We report on the inherent heterogeneity of RBC perfusion within primitive vasculatures, and uncover a strong association between RBC depletion and vessel regression. Our work indicates the essential role of RBC dynamics in the establishment of regional WSS differences driving vascular remodelling. The RBC-driven process of pruning cell-depleted vessels not only importantly contributes to the optimal patterning of vascular networks during development, but also provides a remodelling mechanism to support clinical findings of microangiopathic complications associated with impaired RBC deformability in diseases such as diabetes mellitus and hypertension.

## 1 Introduction

Sprouting angiogenesis is an essential vascularisation mechanism and consists of two well-differentiated phases: sprouting and remodelling (1, 2). During the sprouting phase, a primitive network of vessels is laid out in response to proangiogenic factors *via* a well-established programme of cellular and molecular events (see (3) for a review). The re-modelling phase is responsible for overhauling this primitive network into a hierarchical structure that can efficiently implement the transport capabilities of the cardiovascular system. During the remodelling phase, extensive vessel pruning is achieved primarily *via* dynamic rearrangement of endothelial cells (ECs) (4).

The mechanobiological regulation of ECs has been extensively studied, and it is known that ECs respond to their haemodynamic environment (5). Studies in various animal models have shown that blood flow provides mechanical cues to drive vascular remodelling (*e.g.* chick embryo (6), mouse yolk sac (7, 8), mouse aortic arch (9), zebrafish embryo (10, 11, 12, 4), and mouse retina (4)). Furthermore, these studies have uncovered multiple molecular regulators of EC response to blood shear stress, such as VEGF (13), Wnt (14), Notch (15, 16), TGF*β*/BMP (17, 18, 19), DACH1 (20) and KLF2 (21, 22).

In previous works, we demonstrated that blood shear stress coordinates EC migratory behaviour to achieve vessel regression during the remodelling phase (4, 14, 18). In particular, differences in wall shear stress (WSS) between neighbouring vessel segments lead to polarisation and migration of ECs away from vessel segments experiencing low WSS. In these studies, WSS was calculated using a mathematical flow model that assumes generalised Newtonian rheology (*i.e.* a homogeneous fluid of variable viscosity) without considering the presence of individual red blood cells (RBCs). However, recent computational studies in microscale vessels have demonstrated that RBCs leave transient WSS luminal footprints and therefore could non-trivially modify the local WSS differences driving vascular remodelling (23, 24, 25). This effect of RBCs is closely related to their crucial role in the formation of cell-free layer and the regulation of effective viscosity as well as flow field in microvascular blood flow (26, 27, 28, 29, 30, 31).

In the current study, we propose that the cellular nature of blood (*i.e.* primarily a suspension of RBCs) plays a key role in establishing the local WSS differences driving vascular remodelling. We approach the problem computationally based on the mouse retina model of angiogenesis and provide experimental validation in a developmental zebrafish model. We present simulations of cellular blood flow in vascular networks undergoing remodelling and characterise the bulk flow and RBC dynamics in them. Remarkably, we uncover a previously unreported high-level heterogeneity in RBC perfusion throughout the developing network and a strong association between RBC depletion and vessel regression, which we further confirm experimentally in our zebrafish model *via* live imaging. Furthermore, our experiments confirm previous findings with additional insights that the presence of RBCs is required for effective vascular remodelling at a whole plexus level (7). Finally, we demonstrate that RBC depletion is primarily driven by the plasma skimming effect, *i.e.* the uneven distribution of RBC volume fraction at microvascular bifurcations (32, 33), but uncover important deviations from existing theory caused by asymmetry of the cross-sectional distribution of haematocrit in some feeding vessels.

In line with our previous findings on WSS-modulated EC migration, we argue here that RBC depletion constitutes a mechanism for the enhancement of local WSS differences driving vascular remodelling. This is attributed to the direct relationship between RBC volume fraction, effective viscosity, and WSS. Additionally, we speculate that vascular remodelling driven by the principle of removing RBC-poor vessels from the primitive vasculature will lead to a network layout that avoids portions of the tissue being vascularised but with poorly oxygenated blood. This RBC-driven process, which is highly dynamical and emergent in nature, can importantly contribute to the optimal patterning of vascular networks during development. Conversely, it provides a vascular remodelling mechanism to support recent clinical findings in diabetes mellitus and hypertension (34, 35, 36, 37) reporting that the onset and progression of microangiopathy may arise from altered RBC mechanical properties (*e.g.* impaired deformability) that disrupts the microcirculatory blood flow: a hypothesis first proposed by Simpson in the 1980s (38).

## 2 Results

### 2.1 Simulation of cellular blood flow in microvascular network: validation versus experimental measurements

The vascular plexus of a wild-type mouse retina at postnatal day 5 (P5) was imaged and binarised following staining for collagen IV matrix sleeves (Col.IV) and ICAM2 luminal reporter (see protocol in Sec. 5.1.1). The ICAM2 mask delineates the perfused vessels in the network while the pixel-wise difference between the Col.IV and ICAM2 masks highlights vessel segments undergoing remodelling (Fig. 1a) as demonstrated in Franco *et al.* (4). Therefore, the Col.IV mask constitutes a good approximation of the network morphology prior to these remodelling events. Indeed, the network recapitulates the log-normal distribution of vessel diameters previously reported by Bernabeu *et al.* (41), with a maximum value of 45 *μ*m and a mean of of 11.85 *μ*m (Fig. 1b). Based on our previously proposed approach (42), we construct 3D flow models from the Col.IV binary image and run a whole-plexus blood flow simulation under the assumption of generalised non-Newtonian blood rheology (see the methods in Sec. 5.2.1). This is followed by simulations of RBC flow in designated regions of interest (ROIs) from the capillary bed (*e.g.* ROI-1, ROI-2 and ROI-3 in insets of Fig. 1a) with inflow/outflow boundary conditions obtained from the whole plexus model (see the methods in Sec. 5.2.2). For full details of the computational framework including model configuration and simulation parameters, refer to Fig. S1 and Tables S1–S3 in Sec. S1 of the Supplementary Materials.

**Fig. 1.**
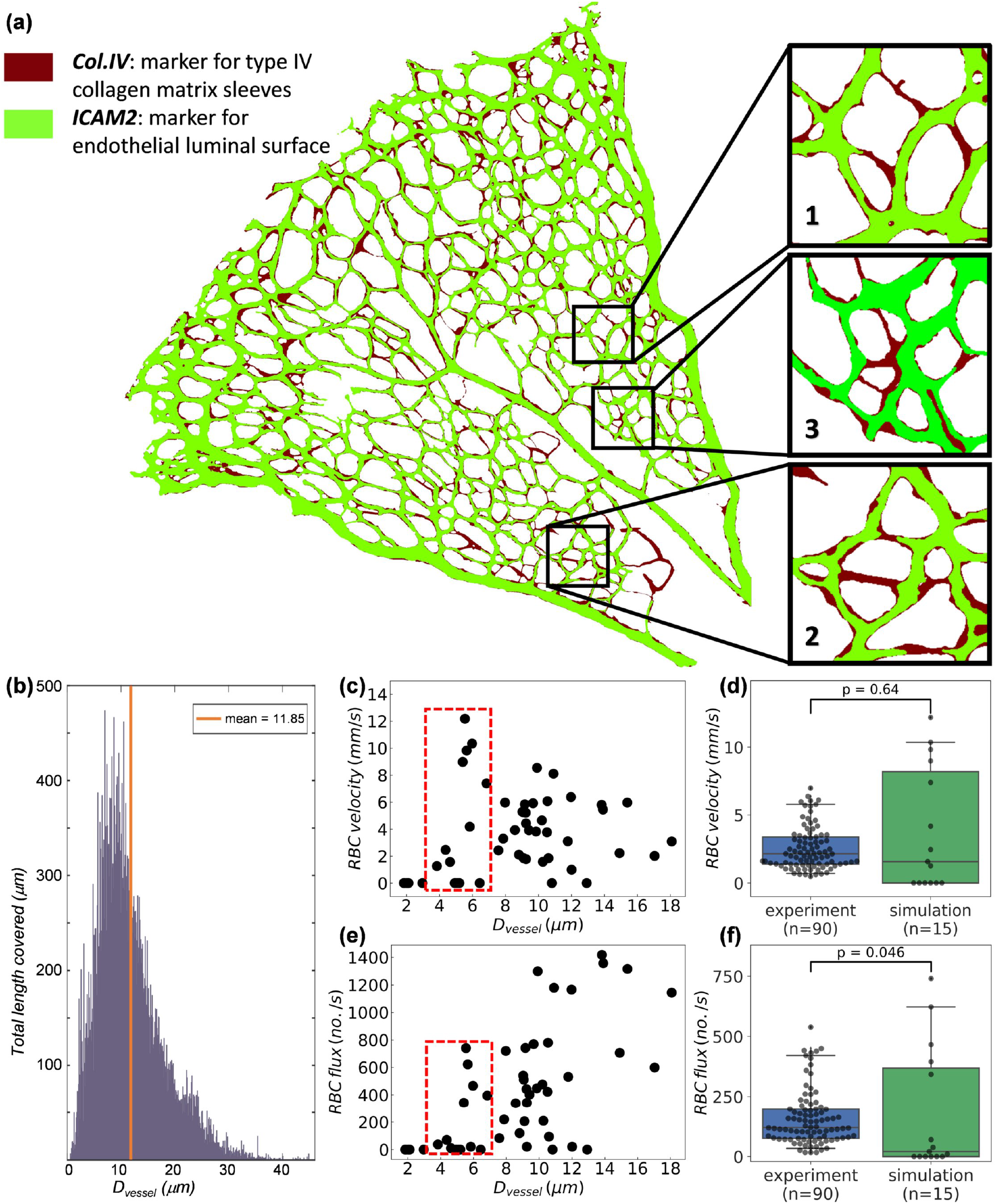
Simulation results of RBC velocity and flux in the primitive vasculature of developing mouse retina. (**a**) A vascular plexus of postnatal day 5 (P5) mouse retina, with vessel lumina and collagen matrix sleeves labelled by ICAM2 (green) and Col.IV (dark red), respectively. The insets show three regions of interest (ROI-1, ROI-2 and ROI-3). (**b**) Network diameter histogram showing the total length covered by vessel segments of certain diameters. (**c**) Simulated RBC velocities and (**e**) RBC fluxes measured in divergent bifurcations within the ROIs. The box of red dashed lines in (c,e) highlight results from small vessels within diameter between 3 and 7 *μ*m. (**d**,**f**) Comparison (Mann-Whitney U test) of simulated RBC velocities and RBC fluxes against *in vivo* measurements (39, 40) from capillary vessels with diameter between 3 and 7 *μ*m.

Our simulation recapitulates volume flow rates in the main artery and veins (running in the axial direction of the retina) of 0.36 *μl*/min and 0.17–0.19 *μl*/min, respectively. These flow rates are in good agreement with *in vivo* measurements recently performed in adult mouse retina of 0.39–0.59 *μl*/min and 0.24 *μL*/min, respectively (39). The RBC velocities and fluxes calculated in the simulated ROIs are in the range 0–12.5 mm/s (Fig. 1c) and 0–1400 RBC/s (Fig. 1e), respectively. These results are validated against *in vivo* single-cell velocimetry data obtained in capillary vessels of diameter 3–7 *μ*m (39, 40). Both the simulation and experimental RBC velocities follow a skewed distribution and show good agreement with median values of 2.14 mm/s and 1.57 mm/s, respectively (*p* ⨠ 0.05, Fig. 1d). The RBC fluxes are significantly higher in experiments than in simulations with median values of 120 RBC/s and 21 RBC/s, respectively (*p* < 0.05, Fig. 1f). The main reason for the lower median observed in simulations is the presence of a number of vessel segments with zero or negligible RBC fluxes (Fig. 1f), which will become the focus of study in the following section Sec. 2.2.

### 2.2 Strong association between RBC depletion and vessel regression in developing retinal vasculature

Having provided evidence of the broad range of RBC fluxes within the ROIs, we now investigate a potential association between the RBC dynamics and vessel regression. We classify ROI vessel segments into three groups (Fig. 2a): the “lumenised” group shows positive signals in both Col.IV and ICAM2, featuring open lumina; the “stenosis” group shows positive signal in Col.IV and partial-positive/partial-negative signal in ICAM2, featuring vessel stenosis (an early stage of vessel regression (4)); the “regression” group shows positive signal in Col.IV and negative signal in ICAM2, featuring vessel regression.

**Fig. 2.**
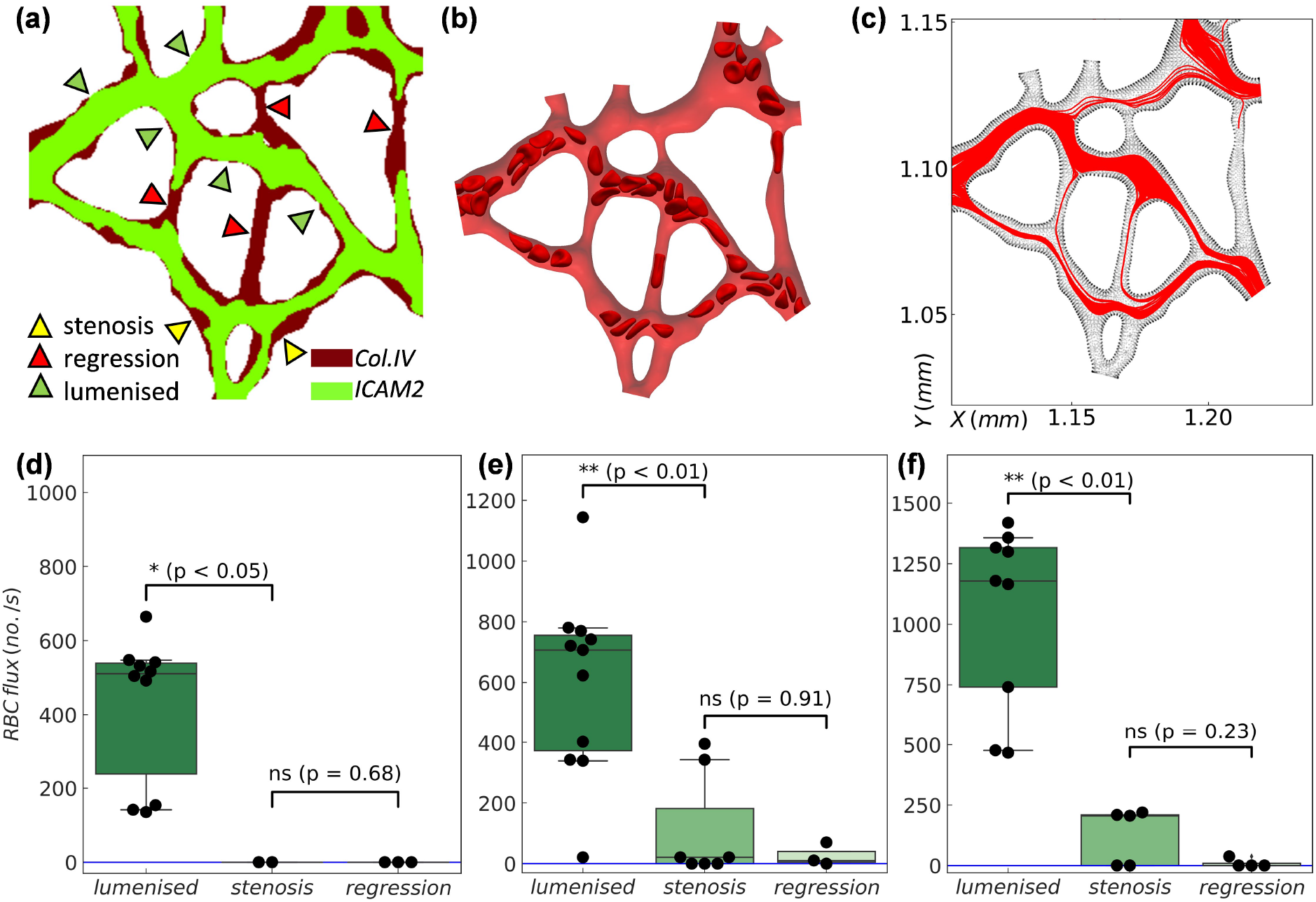
Association between RBC depletion and vessel regression in the developing retinal network. (**a**) Characterisation of vessel segments based on ICAM2 and Col.IV signals (here showing ROI-2 as in Fig. 1). The investigated vessels are divided into three groups, *i.e.* lumenised, regression and stenosis. (**b**) Corresponding simulation of the cellular blood flow in ROI-2. (**c**) Combined cell trajectories over time in ROI-2 through-out the simulation. (**d**–**f**) Quantification of time-average RBC fluxes in ROI-1, ROI-2 and ROI-3, respectively. The statistical analysis is performed using Mann–Whitney U test. The sample sizes for the three groups are *n* = 10, 2, 3 in (d); *n* = 11, 7, 3 in (e); *n* = 9, 5, 4 in (f).

Further examination of our cellular flow simulations reveals that most RBC-depleted vessels in the simulation coincide with vessels in the regressed group (Fig. 2a-b). To quantify this, we record the trajectories of all RBCs within each ROI throughout the simulation and study their density across the ROIs (Fig. 2c and Fig. S2 in the Supplementary Materials). In general, a vessel segment with higher density represents good RBC perfusion, whereas those with low density indicate poor RBC perfusion, or RBC depletion. Subsequently, the time-averaged RBC flux within each vessel segment is calculated and assigned to the three groups under study (Fig. 2d-f). Our analysis demonstrates that vessel segments in the lumenised group have significantly higher RBC fluxes than those in the stenosis group and the regression group (*p* < 0.05 for ROI-1 and *p* < 0.01 for ROI-2 and ROI-3, Fig. 2d-f). Meanwhile, the difference in RBC flux between the stenosis group and the regression group is not significant (*p* ⨠ 0.05 for all ROIs, Fig. 2d-f). These results support a strong association between RBC depletion and vessel regression.

### 2.3 *In vivo* validation of the effect of RBC perfusion on vascular remodelling in zebrafish caudal vein plexus

To provide experimental confirmation of the association between RBC depletion and vessel regression predicted by our computational model, we turned to the zebrafish model of vascular development, where simultaneous live imaging of vessel remodelling and RBC dynamics is possible. We chose the caudal vein plexus (CVP) for observation 48–72 hours post fertilisation (hpf), a period during which gradual remodelling of the plexus down to a single, well-defined vascular tube begins (43). Comparison was made between control (ctl) morpholino oligomer (MO) fish with normal RBC perfusion (Fig. 3a) and gata1 MO fish not carrying RBCs in the bloodstream (Fig. 3b, see Sec. 5.1.2 for experimental details). Our time-lapse imaging of the CVP in ctl MO fish captures heterogeneous RBC perfusion (Fig. 3c) leading to multiple findings of intermittent and complete RBC depletion in vessel segments (Fig. 3d) followed by vessel stenosis (Fig. 3e) and eventual regression (Fig. 3f). These *in vivo* findings therefore confirm our computational predictions in Sec. 2.2.

**Fig. 3.**
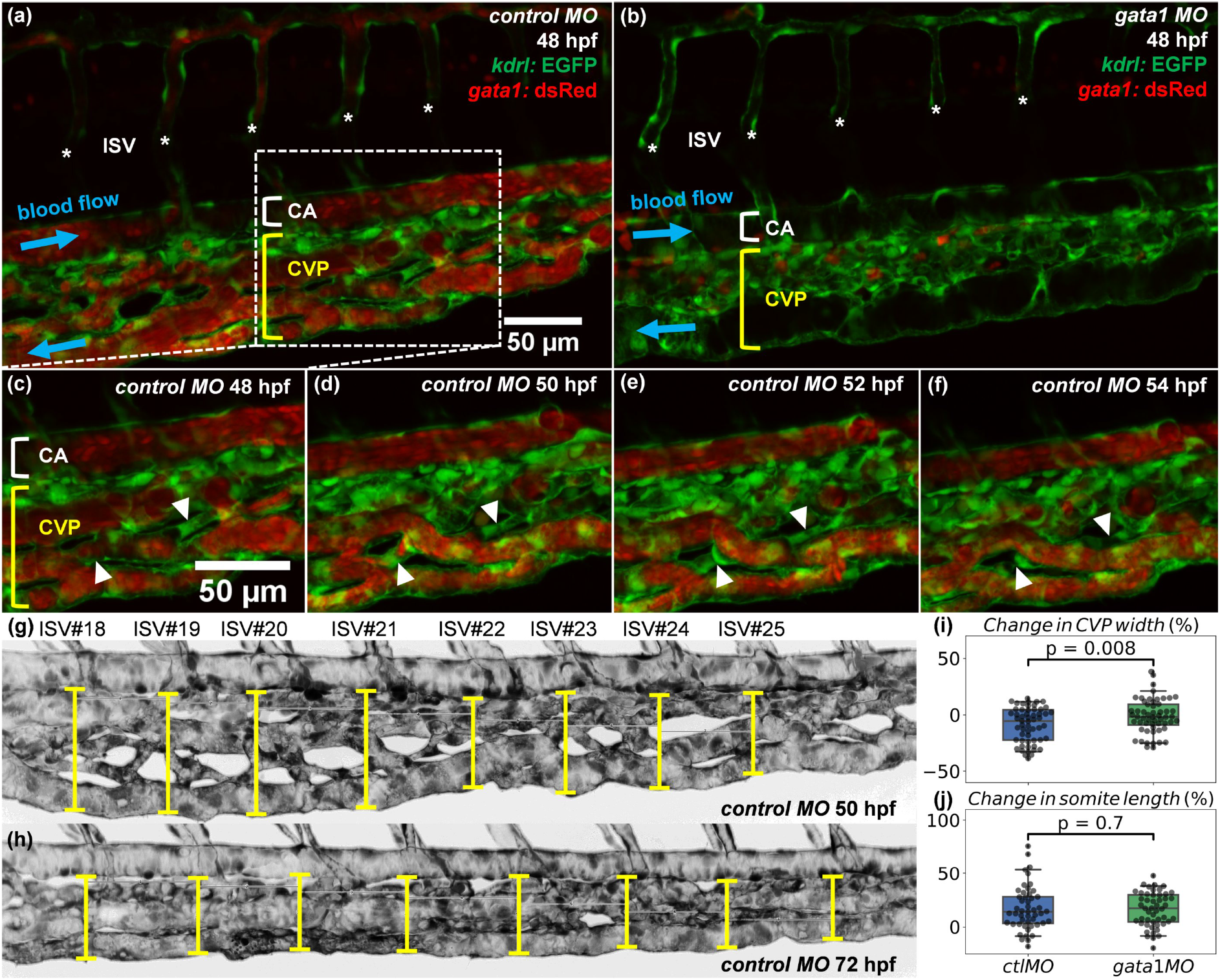
Time-lapse imaging of RBC perfusion and vascular remodelling in zebrafish caudal vein plexus. (**a**,**b**) Two exemplar caudal vein plexuses (CVPs, indicated by a square bracket in yellow) from a 48 hpf ctl MO embryo (with RBC perfusion) and a 48 hpf gata1 MO embryo (*Tg(GATA-1:eGFP)*, without RBC perfusion). The intersegmental vessels (ISVs) are marked with asterisks, and the caudal artery (CA) is indicated by square bracket in white. The RBC precursors in (b) are located outside the vasculature and not circulating within the blood stream. (**c**–**f**) Time sequence showing vessel regression events in a region of interest extracted from the ctl MO embryo in panel (a), where two vessel segments marked by white triangles are pruned over time (*t* = 48 hpf, 50 hpf, 52 hpf, 54 hpf). (**g**,**h**) Exemplar measurements of CVP widths (indicated by capped lines) at *t* = 50 hpf and *t* = 72 hpf along the anterior-posterior axis of a ctl MO embryo (Z-projection image) at positions given by eight consecutive ISVs (ISV 18–ISV 25). The length of corresponding somites (seven counted here for each fish) are estimated by the inter-ISV distances. (**i**) Normalised CVP width change and (**j**) normalised somite length change at *t* = 72 hpf against *t* = 50 hpf (relative change in percentages), calculated from measurements of the ctl MO group and the gata1 MO group (each containing 7 embryos). The statistical analysis in (i,j) is performed using Welch’s T test, with *n* = 56 in (i) and *n* = 49 in (j).

Next, we investigated how RBC deletion in gata1 MO fish impacts CVP remodelling at a network level. We measured CVP widths at standardised locations along the anterior-posterior fish axis given by the positions of eight consecutive intersegmental vessels (ISVs), for both the ctl MO and gata1 MO groups (7 fishes in each group) at 50 hpf and 72 hpf (see Fig. 3g-h for an example). During this period of time, substantial remodelling leading to narrowing of the plexus is observed in the wild type (agreeing with (43)). Furthermore, significantly larger reduction of CVP width in the ctl MO group is found in comparison to the gata1 MO group, accounting for 9% and 1% of relative width change (mean reductions 7.64 *μ*m and 2.01 *μ*m), respectively (*p* < 0.01, Fig. 3i). To exclude that differences in fish growth may confound these results, the longitudinal extension of the somites is also examined, for which no significant difference is found between the ctl MO and gata1 MO groups (*p* ⨠ 0.05, Fig. 3j). Taken together, this several-fold difference in CVP width change implies that the presence of RBCs is necessary for normal CVP remodelling 50–72 hpf, and therefore the heterogeneity in RBC perfusion described in Fig. 3c-f does play a role in orchestrating network-level remodelling.

### 2.4 Network RBC depletion not predictable by vessel diameter

The absence of RBCs in some vessel segments in both our simulations and experiments poses a crucial question about cellular flow in developmental vascular network: what is the governing mechanism that determines which vessels to be perfused with cells and which to be devoid of? It is tempting to speculate that the pattern of vessel RBC depletion is merely a size-exclusion effect; namely, certain vessel segments are simply too narrow to allow cells to pass through. However, the vessel diameters encountered in the present simulations (about 2–16 *μ*m) are unlikely to be the dominant factor affecting RBC transit, since the high deformability of RBCs under physiological conditions enables them to pass through exceedingly small passages as narrow as 1–2 *μ*m (44, 45, 46). Indeed, the material model adopted for the RBC membrane allows the cells to deform and adopt highly elongated shapes to flow through all narrow capillaries that they enter in our simulations (with one exception only), whereas some larger vessels are devoid of RBCs (see Movies S1-S3 of the Supplementary Materials).

Furthermore, we evaluate the degree of RBC enrichment/depletion by examining the sign and magnitude of 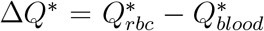 (positive for enrichment and negative for depletion, see definitions in Sec. 5.3.1) against vessel diameter *D_vessel_* for all child branches studied here (44 in total, Fig. 4a). RBC depletion occurs throughout the whole range of vessel sizes investigated (*D*_*vessel*_ ∈ [2, 16] *μ*m). Notably, for one child branch with *D*_*vessel*_ ≈ 9 *μ*m (larger than the physiological RBC diameter), Δ*Q*\* is *−*0.2, indicating a 20% reduction in RBC transit. Based on these findings, we conclude that RBC depletion is not a size-exclusion effect. Meanwhile, it is found that RBC enrichment happens only in medium/large vessels (*D*_*vessel*_ > 5 *μ*m). Within the intermediate diameter range *D*_*vessel*_ ∈ [5, 12] *μ*m, the vessels have nearly equal chances of being enriched or depleted.

**Fig. 4.**
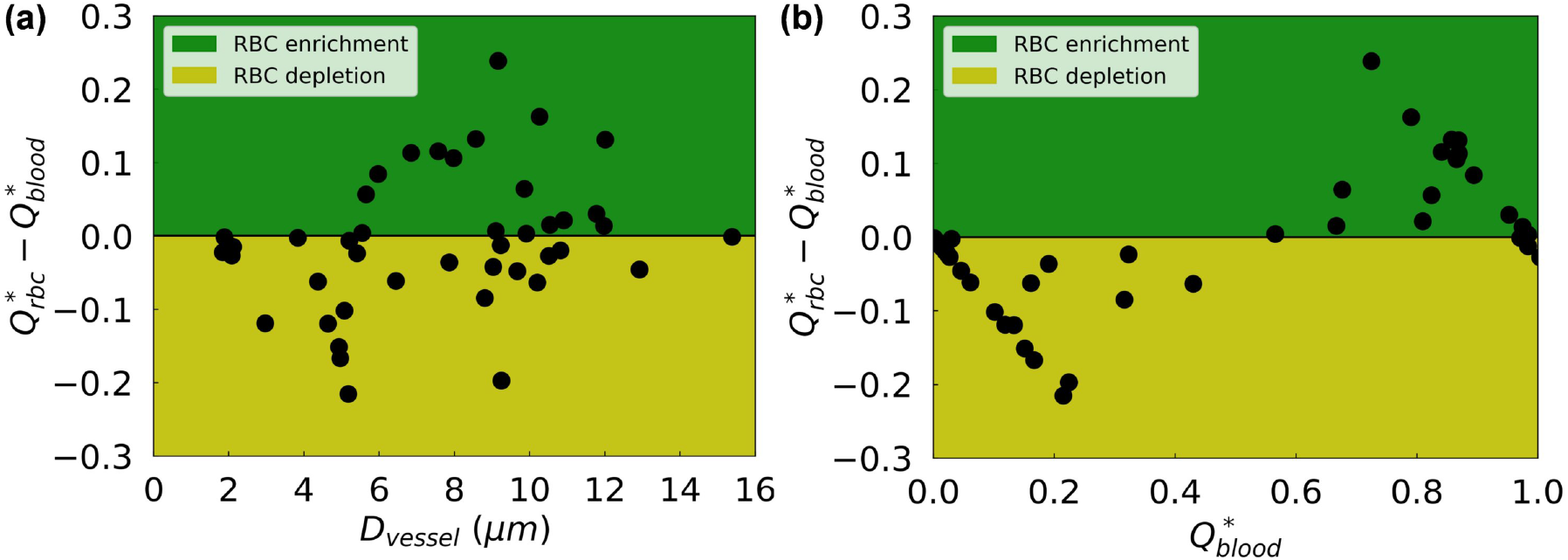
Quantification of RBC depletion in the developing retinal network. 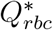 and 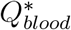 represent the normalised RBC flux and normalised blood flow in a given child vessel (with diameter *D*_*vessel*_) relative to those in its parent vessel, respectively. The variable 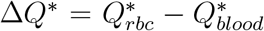 serves as a disproportionality index of flow-mediated RBC partitioning, based on the sign of which the vessels are classified as “RBC-depletion” (negative Δ*Q*\*, yellow patch) and “RBC-enrichment” (positive Δ*Q*\*, green patch). The disproportionality indices for all investigated vessel segments are sorted against (**a**) vessel diameter *D_vessel_* and (**b**) normalised blood flow 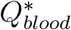, respectively. The analysed vessel segments in this plot are extracted from the three ROIs in Fig. 1.

### 2.5 Plasma skimming as a mechanism for RBC depletion in developing vascular network

Having demonstrated that size-exclusion effect is not sufficient to explain the RBC depletion observed in Sec. 2.2, we now turn our attention to potential haemodynamic mechanisms. First, we observe that characterising 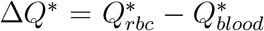 against the haemo-dynamic indicator 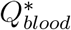 does satisfactorily separate the RBC-depletion zone from the RBC-enrichment zone (Fig. 4b). This finding is in line with the plasma skimming effect described by empirical models such as the widely-employed phase-separation model (PSM, see introduction in Sec. 5.3.2), first proposed by Pries and co-workers (47, 48).

To qualitatively assess the agreement between our data and the PSM, we plot the fractional RBC fluxes 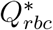 in individual child branches (44 studied here) of any divergent bifurcation against their fractional blood flow 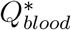 (Fig. 5a). The distribution is indeed reminiscent of the sigmoidal relationship predicted by the PSM, where 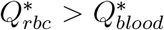 for 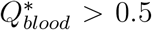 up to a threshold where 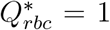 (and the opposite effect for 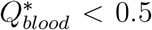 down to 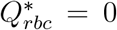). Furthermore, albeit with occasional exceptions, the child branch relatively smaller in size (red circles) within a bifurcation tends to receive lower blood flow and consequently fewer RBCs, whereas the larger child branch (blue squares) is more likely to have higher blood flow and attract more RBCs (inset of Fig. 5a). Note that “smaller” or “larger” here is a relative notation between the two child branches within a bifurcation, instead of a measure of the absolute vessel size.

**Fig. 5.**
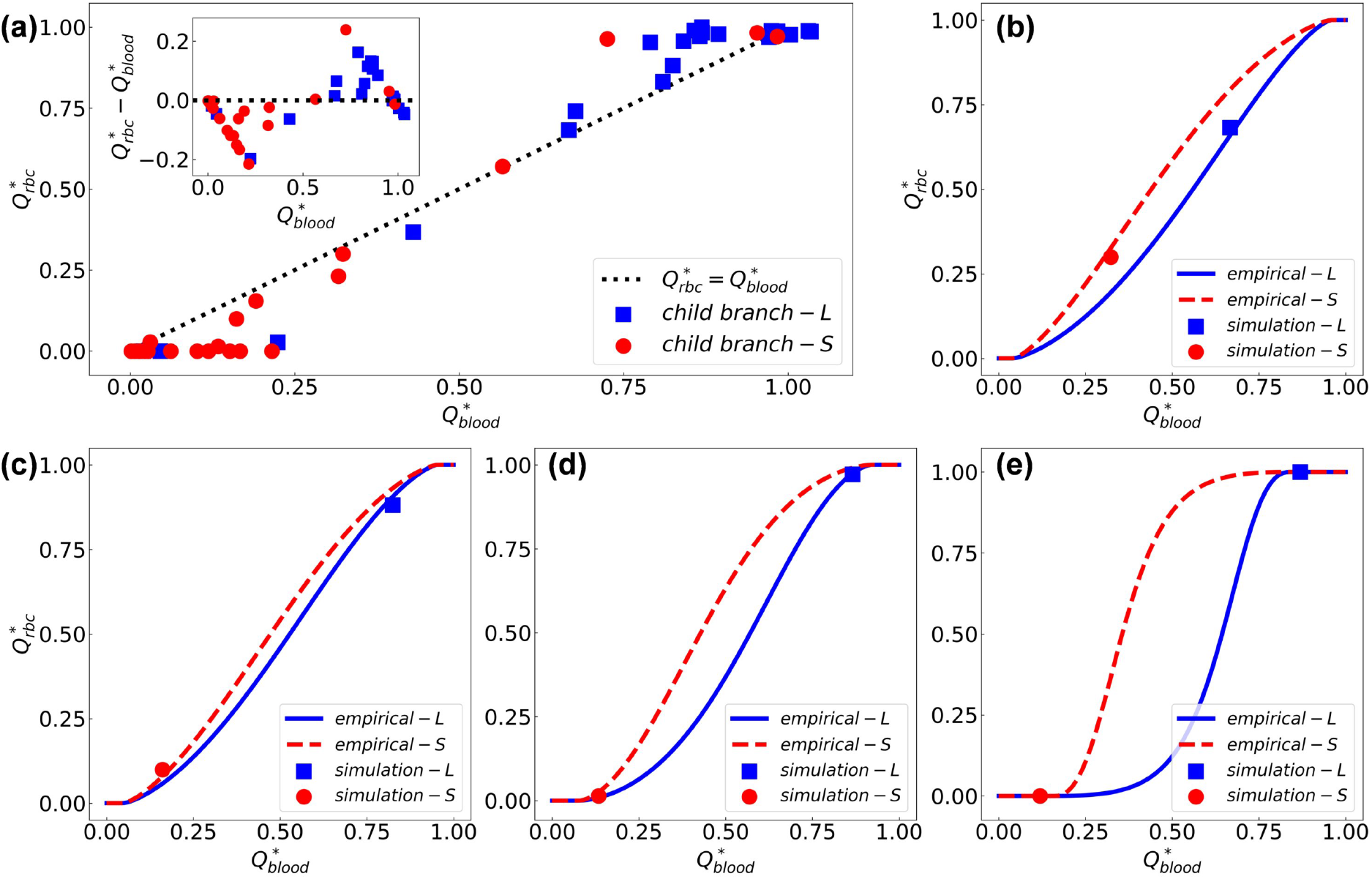
Comparison of simulation data with empirical predictions by the phase separation model (49). (**a**) Simulation data of fractional RBC flux 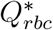 against fractional blood flow 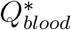 in the relatively larger child branch “L” (blue squares) and smaller child branch “S” (red circles) from all investigated bifurcations. The inset shows similar results as in Fig. 4b (characterising the disproportionality index 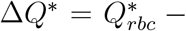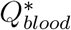 against 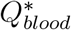), but with additional information of relative vessel size for child branches in each bifurcation. The black dotted line represents a linear hypothesis for 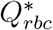 and 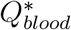 in the absence of plasma skimming. (**b**–**e**) Four exemplar bifurcations in which the simulation data (squares and circles) agree well with empirical predictions (solid lines) for both the “L” and “S” child branches.

Furthermore, we compare our simulation data at each bifurcation with the prediction given by the PSM model (Equations (1)–(4)). We observe errors of less than 5% for 18 out of 22 bifurcations (see Fig.s S3–S4 for individual comparison and Table S4 for complete error evaluation in Sec. S2 of the Supplementary Materials). Fig. 5b–e demonstrate four cases of good agreement. In the first bifurcation, both child branches have considerable proportions of blood flow (roughly 30% and 70%, respectively) and are well-perfused by RBCs, with the fractional RBC fluxes matching the PSM predictions (Fig. 5b). In the second bifurcation, most RBCs 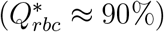 enter the relatively larger child branch as it receives more than 80% of the blood flow from the parent branch (Fig. 5c). In the third and fourth bifurcations (Fig. 5d–e), the smaller child branch is nearly devoid of cells as the relatively larger branch attracts almost all RBCs from the feeding vessel owing to its predominantly higher proportion of blood flow 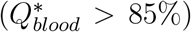. Larger than 5% deviations from the PSM are observed in the remaining 4 of the 22 bifurcations studied.

Shown in Fig. 6a–b are two cases where the simulated RBC fluxes in the bifurcation deviate from the empirically predicted values. Interestingly, the RBC flux contrast between the two child branches is slightly underestimated by the PSM model in the first case (Fig. 6a) whereas it is substantially overestimated in the second case (Fig. 6b). To investigate these disagreements, we examine the flow streamlines in the mid-plane of the bifurcation and calculate the relative distance *χ* of the stagnation streamline separating the flow entering the left branch from that entering the right branch (Fig. 6c–d). Clearly, the higher-flow branch (child branch with larger 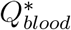) receives proportionally more streamlines from the parent vessel and should therefore receive more RBCs accordingly provided that the cells are axisymmetrically distributed in the parent branch (as assumed by the empirical model of PSM).

**Fig. 6.**
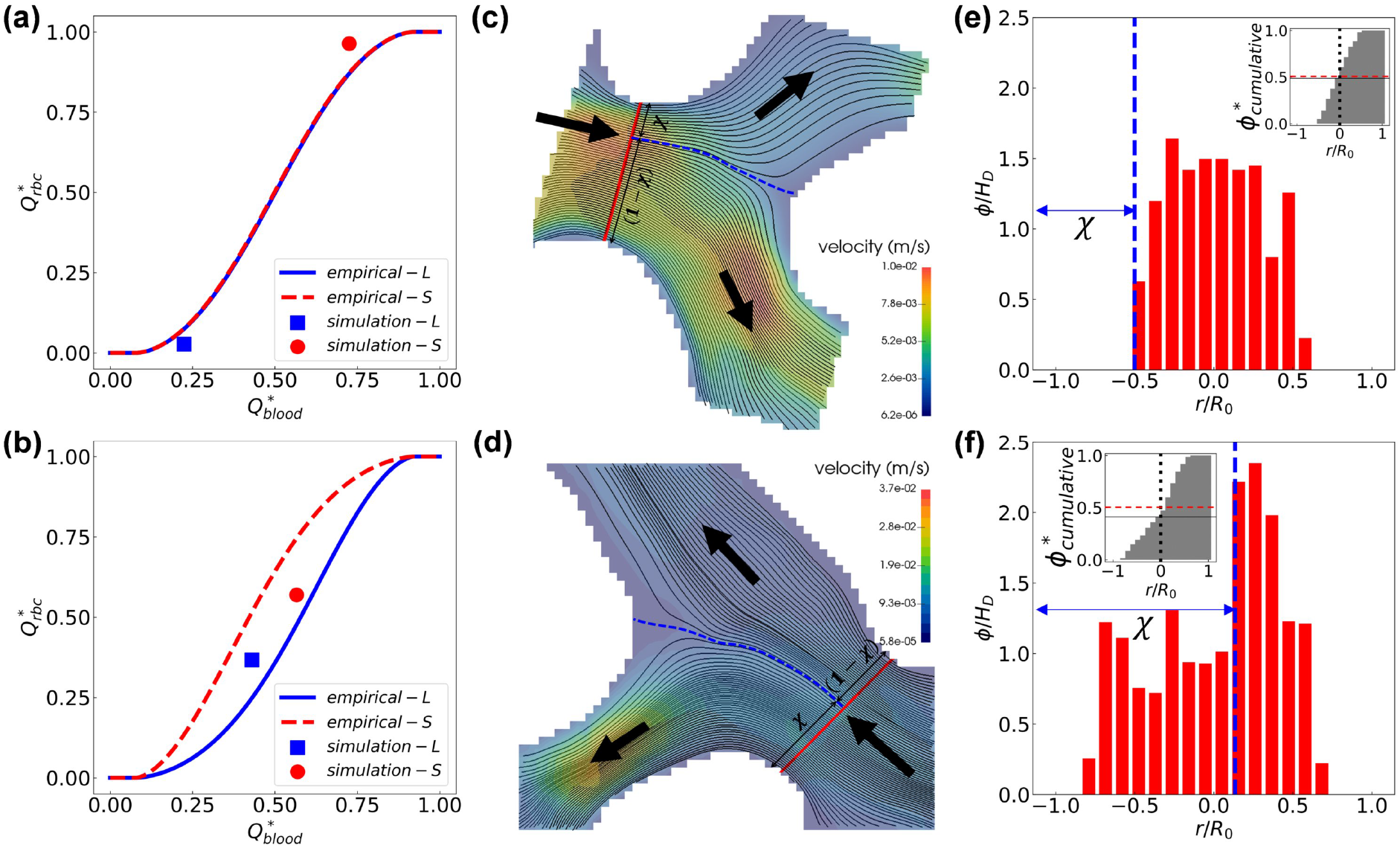
Occasional deviation of simulation data from the empirical model (49) due to the asymmetry of haematocrit profile in the parent branch. (**a**–**b**) Two exemplar divergent bifurcations for which the simulation data (circle dots) deviate from PSM predictions (solid lines). (**c**–**d**) Visualisation of the flow streamlines separated into the child branches on the mid-plane of the bifurcation (extracted from the 3D simulation). The blue dashed line indicates the location of the separation surface. (**e**–**f**) Cross-sectional haematocrit profile in the parent branch, at a position marked by the red solid line in (c–d). The blue dashed line corresponds to the separation surface of flow streamlines as in (c–d). The insets of (e–f) show the cumulative haematocrit distribution corresponding to the haematocrit profile.

Next, we investigate whether the assumption of haematocrit axisymmetry by the PSM holds in our simulations. To this end, we calculate the cross-sectional distribution of RBCs in the parent branch. We find clearly asymmetric distributions, especially for case two (Fig. 6e–f). In both cases, the haematocrit profile is skewed towards the right-hand side of the bifurcation, with a cumulative haematocrit to the left-hand side of the vessel centreline either slightly or substantially smaller than 0.5 (see insets of Fig. 6e–f). Such haematocrit asymmetry makes the downstream child branch on the right-hand side inherently advantageous for RBC intake (hereafter referred to as haematocrit-favoured branch). In case one, the higher-flow branch coincides with the haematocrit-favoured branch, resulting in enhanced RBC flux difference between the two child branches; whereas in case two, the higher-flow branch differs from the haematocrit-favoured branch, thus attenuating the RBC flux difference. A full description of the mechanism leading to this skewness, which originates from the interplay between the complex geometry and the emerging RBC behaviour in the microvasculature, is out of the scope of the present study and will be explored in future work.

## 3 Discussion

Earlier studies have extensively explored the effect of blood flow on cardiovascular development and it is well accepted that haemodynamic cues are essential for vascular development (7, 11, 12). Previous work by our groups further established the pivotal role of regional WSS differences, which were found to locally modulate the polarised migration of ECs into high-shear vessel segments and cause the regression of adjacent segments experiencing low shear (4, 14, 18). However, because the quantification of WSS in these studies relied on mathematical flow models assuming simplified blood rheology, the question of how the presence of RBCs and their profound impact on microvascular haemodynamics affects vascular patterning has not been addressed. Recent simulations of cellular blood flow in single microvessels have shown that RBCs can non-trivially modify both the mean and oscillatory components of local WSS (23, 25), thereby suggesting a salient impact of RBCs on the mechanotransduction of fluid forces during angiogenesis. Based on these findings, we hypothesise that RBCs in the developing mouse retina play an active role in the course of vascular patterning towards a functional network *via* altering WSS spatially and temporally.

In the current study, we simulated RBC dynamics in the vascular plexus of a wild-type mouse retina at P5. The computed RBC velocities are in good agreement with *in vivo* measurements made in adult mouse retinas (39, 40). However, differences arise in terms of RBC fluxes (more heterogeneous distribution with lower median value, Fig. 1f). We attribute this discrepancy to structural differences between the developmental stage simulated and the adult stage considered for validation. We hypothesise that as the primitive network remodels, the simulated RBC fluxes will become closer to the *in vivo* measurements in adult mice. Future advances in live imaging of the mouse retina model will contribute to elucidating this process.

There are other sources of uncertainty in the current simulations remaining to be quantified. First, the discharge haematocrit at the inlets of all ROIs was fixed at 20% for consistency. This value is in line with the average microvascular haematocrit level reported in the literature, *i.e.* 23% ± 14% by (50), but may not necessarily reflect the haematocrit levels of individual mice studied in (39, 40) and used for validation in our study. Second, the flow in realistic microvessels with smaller lumen size than the RBC diameter is typically impeded due to the close contact of RBCs with the endothelial surface layer (ESL, consisting of glycocalyx and/or cilia), whereas in our present *in silico* model such microscopic structures on the vessel wall are not considered. Prior studies in glass tubes where the ESL was absent demonstrated a two-fold decrease in flow resistance when compared to *in vivo* conditions (51, 52).

Our simulations of cellular blood flow reveal a previously unreported high-level heterogeneity in RBC perfusion throughout the developing vascular network, where a number of plasma vessels exist with rare RBC transit over time, and a strong association between RBC depletion and vessel regression. From a mechanistic point of view, the effective viscosity in neighbouring RBC-depleted and RBC-enriched branches can differ substantially. Such disparity in viscosity will enhance the regional WSS difference between neighbouring branches, which can in turn promote the pruning of the vessel segment depleted of RBCs according to our previously reported mechanisms (4).

We provide further experimental confirmation of the above findings in a developmental zebrafish model, which is amenable to live imaging. In agreement with Lucitti *et al.* (7), we show that the presence of RBCs is necessary for effective remodelling at a whole plexus level. Furthermore, we extend the conceptual model by demonstrating that intermittent and complete RBC depletion selects vessels for pruning, which are likely to experience low shear and become unfavoured during vascular remodelling. Apart from their impact on the WSS differences, Xiong *et al.* (53) recently reported that the signalling role of RBCs also influences vessel remodelling as erythrocyte-specific sphingosine-1-phosphate was found crucial for vascular stabilisation and maturation during embryonic development. Understanding the interplay between haemodynamic forces and secreted factors derived from RBCs, however, is beyond the scope of the present study.

To fully understand the perfusion of RBCs within the developing mouse retina, we asked the question: what is the mechanism behind RBC depletion/enrichment in the primitive network? Quantification of RBC fluxes against vessel diameter and blood flow in individual bifurcations rules out vessel-size exclusion as the primary factor. Instead, the empirical model by Pries *et al.* (47, 49) based on the plasma skimming effect explains the uneven partitioning of RBC fluxes satisfactorily in 18 out of 22 cases, therefore implying that the distribution of RBCs within the developing network is flow-mediated rather than geometry-dominant.

For the 4 cases where predictions of the empirical model do not satisfactorily match the simulation data, we find considerable haematocrit asymmetry in the cross-sections of the feeding branches, which is against the central assumption of axisymmetric haematocrit profile by the model. This makes accurate prediction of the RBC perfusion in a given vascular network challenging without certain knowledge of cross-sectional cell distributions. Our observation of haematocrit-favoured and haematocrit-unfavoured vessel branches at microvascular bifurcations is in line with recent *in vitro* findings of reversal of the classic haematocrit partitioning which otherwise always favours the higher-flow branch (54, 55, 56, 57). Similar reverse partitioning was reported by simulations of cellular blood flow in microvascular networks with vessel diameters designed following Horton’s law (30). Our group has recently identified reduced interbifurcation distance and complex branching topology as sources of haematocrit asymmetry in the context of tumour blood flow. Furthermore, we propose a role for the resulting abnormal partitioning in establishing tumour tissue hypoxia (58).

The asymmetry not only applies to the haematocrit profile, but also the velocity profile. For a majority of microcirculatory models, the Poiseuille law has been employed to simplify the haemodynamics and reduce computational cost, manifested by parabolic velocity profiles featuring a peak velocity at the centreline of microvessels. Our results counter such a simplification, as the velocity profile in a capillary vessel can significantly deviate from a parabola and become skewed over time in the presence of travelling RBCs (Fig. S5 in Sec. S3 of the Supplementary Materials). Similar deviation of velocity profiles has also been confirmed by imaging data from living mouse retina, where a 39% error in flow estimation was reported if assuming a parabolic profile (39).

Finally, the implications of the uncovered association between RBC depletion and vessel regression are not limited to developmental vascular remodelling. Diseases such as diabetes mellitus and hypertension lead to changes in RBC deformability (59, 60, 61) and this have been associated with vascular complications such as diabetic retinopathy or nephropathy in several clinical studies (34, 35, 36, 37). However, the magnitude of the changes in RBC deformability reported in the literature (ranging between 10%-50% percent (62, 63, 37) to several folds (64, 65)) would not support a model where such biomechanical changes alone are sufficient to cause vessel occlusion since Freund (44) showed that order of magnitude changes in shear modulus are required to impede RBC transit through narrow passages. In contrast, our findings support a new concept where changes in the mechanical properties of the RBC membrane leading to abnormal haematocrit partitioning at bifurcations (*via* altered radial distributions of RBCs (66, 67, 68)) would reintroduce, in adult networks, the differences in WSS driving developmental vascular remodelling. Future work should investigate whether these WSS differences are sufficient to trigger pathological vascular remodelling.

## 4 Concluding remarks

In summary, our study reports a new mechanism for enhancement of the WSS differences driving vascular remodelling during development. These enhanced differences arise due to the highly heterogeneous distribution of RBCs within the primitive plexus, which is primarily governed by the plasma skimming effect. Additionally, we speculate that vascular remodelling driven by the principle of removing RBC-poor vessels from the primitive vasculature will lead to a network layout that avoids portions of the tissue being vascularised but poorly oxygenated. This RBC-driven process, which is highly dynamical and emerging in nature, can importantly contribute to the optimal patterning of vascular networks during development. Conversely, it provides a vascular remodelling mechanism capable of linking changes in RBC deformability reported in diseases such as diabetes mellitus or hypertension and the associated microangiopathic complications. Beyond these findings, our study also has important implications for the mathematical modelling of microvascular haemodynamics in general. In a network of microvessels, multiple effects inexplicable by continuum flow models (albeit widely employed for modelling blood flow in complex vascular networks by existing studies) occur owing to the particulate nature of blood, *i.e.* essentially a suspension of RBCs. Conventional assumptions such as Poiseuille law, velocity-/haematocrit-profile symmetry, spatial-/time-average accuracy are all subject to scrutiny when quantifying flow variables such as effective viscosity and WSS in the microcirculation, especially for vessels with a lumen smaller than the undeformed size of an RBC (*i.e.* 6.2–8.2 *μ*m).

## 5 Materials and methods

### 5.1 Mouse and zebrafish experiments

#### 5.1.1 Preparation of mouse retina for binary mask acquisition

The mouse strain used in the present study was C57/BL6J. Mice were maintained at the Max Delbrück Center for Molecular Medicine under standard husbandry conditions. Animal procedures were performed in accordance with the animal license X9005/15. Mouse eyes were collected at P5 and fixed with 4% PFA in PBS for 1 h at 4°C, and retinas were then dissected in PBS. Blocking/permeabilisation was performed using Claudio’s Blocking Buffer (CBB, 69), consisting of 1% FBS (Gibco), 3% BSA (Sigma-Aldrich), 0.5% Triton X-100 (Sigma-Aldrich), 0.01% sodium deoxycholate (Sigma-Aldrich), and 0.02% sodium azide (Sigma-Aldrich) in PBS at pH 7.4 for 2 h with rocking at 4°C. Primary antibodies were incubated at the desired concentration in 1:1 CBB/PBS with rocking at 4°C overnight and secondary antibodies were incubated at the desired concentration in 1:1 CBB/PBS for 2 h at room temperature. Retinas were mounted on slides using Vectashield mounting medium (H-1000; Vector Labs).

The following primary and secondary antibodies were used *in vivo*: collagen IV (ref 2150-1470, rabbit; 1:400; AbD Serotec) and ICAM2 (ref 553326, rat; 1:200; BD Biosciences), anti-Rat Alexa 488 (ref A21208, donkey 1:400, Invitrogen), anti-Rabbit Alexa 568 (ref A10042, donkey 1:400, Invitrogen). Complete high-resolution three-dimensional rendering of whole mount retinas were taken using a LSM 780 inverted microscope (Zeiss) equipped with a Plan-Apochromat 63×/1.4 NA DIC objective. Images were taken at room temperature using Zen 2.3 software (Zeiss). Tiled scans of whole retinas were analysed with ImageJ to generate binary masks of ICAM2 and Collagen IV.

#### 5.1.2 Morpholino oligomers, zebrafish husbandry and imaging

Morpholino oligomer (MO) against *gata1* Morpholino (gata1 MO) as described in (70) (sequence 5’-CTGCAAGTGTAGTATTGAAGATGTC-3’) was injected at 8 ng/embryo following (71). A control MO (ctl MO) served the standard control Morpholino with the sequence 5’-CCTCTTACCTCAGTTACAATTTATA-3’ targeting a human beta-globin intron mutation. The control was injected at similar amount of 8 ng/embryo. Zebrafish (*Danio rerio*) were raised and staged as previously described in (72). For growing and breeding of transgenic lines, we complied with regulations of the animal ethics committee at the MDC Berlin (73).

Embryos were anaesthetised in 0.014% tricaine (Tricaine Pharmaq 1000mg/g, PHARMAQ Limited), mounted in plastic petri dishes (94×16 mm - Sarstedt Ref#82.1473) containing 0.014% tricaine, and bathed in E3 media containing 0.007 (0.5x) tricaine and 0.003% PTU. Imaging was performed on an upright 3i spinning-disc confocal microscope using Zeiss Plan-Apochromat 20x/1.0 NA water-dipping objectives. Screening of embryos was performed using a Leica M205 FA stereomicroscope with filter set ET GFP M205FA/M165FC.

### 5.2 Numerical simulation

#### 5.2.1 Whole-plexus simulation

A three-dimensional (3D) flow model of the luminal surface (Fig. S1b) is reconstructed from the Col.IV binary mask (Fig. S1a) using the open-source software *PolNet* (42), under the assumption of circular vessel cross-sections. The flow domain is then discretised into uniform cubic lattice grids with a size of Δ*x*, whose value is chosen to be Δ*x* = 0.5 *μ*m for the mouse retina used here such that the flow can be accurately solved throughout the reconstructed network for reliable results of flow rates and wall shear stresses even in the smallest lumenised vessels.

The whole-plexus simulation in the reconstructed network adopts the non-Newtonian Carreau-Yasuda rheology model (NNCY, 74) following our previous approach (41), where blood is modelled as a generalised Newtonian fluid (homogeneous shear-thinning liquid). By imposing a physiological ocular perfusion pressure (OPP) between the arteriole and venules, a steady flow within the vascular plexus is solved (Fig. S1c) to provide boundary conditions for subsequent RBC simulations in designated regions of interest (ROIs) of the plexus. OPP = 55 mmHg is chosen for the present simulation based on a literature survey and sensitivity analysis conducted in (41). Table S1 provides key parameters of the whole-plexus simulation. The computational framework based on immersed-boundary-lattice-Boltzmann method is detailed in Sec. S1 of the Supplementary Materials.

#### 5.2.2 RBC simulation in network subsets

To create RBC simulations in ROIs of the retinal network where evident vessel regression events are observed (*e.g.* Fig. S1d), we first clip the designated ROI from the whole-plexus as a geometric subset (Fig. S1e). For a ROI subset with *N* open boundaries, we set up (*N* – 1) Poiseuille velocity inlets/outlets (where parabolic velocity profiles are imposed with a centreline velocity of *û*) and one pressure outlet (where a reference pressure *p*_*out*_ = 0 is set). The velocity boundary conditions are obtained from the NNCY simulation through measuring local flow rates *Q* in vessels of interest and subsequently calculating *û* from *Q* under the assumption of Poiseuille flow, following 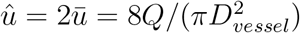. With all the above boundary conditions set (see Table S3), a plasma flow simulation is initiated in the ROI with the fluid viscosity equal to that of plasma *η*_*plasma*_. Once steady flow is achieved and the velocity field is verified against the NNCY simulation, we populate the ROI with RBCs which are continuously fed at a discharge haematocrit of 20 % from all inlets of the subset network (Fig. S1f). Details of the RBC model can be found in Table S2 of the Supplementary Materials.

### 5.3 Data analysis

#### 5.3.1 Vessel selection and quantification of RBC flow

For the study of RBC perfusion within the ROI subsets, we locate all divergent bifurcations (*i.e.* composed of one parent branch and two downstream child branches) encountered by the cellular flow in each ROI *via* exhaustively examining the flow directions (identical between the whole-plexus simulation and the ROI plasma flow simulation) in every single vessels segment (Fig. S2a–c). Only vessel branches from these divergent bifurcations are selected for statistical analysis to ensure independent representation of the systematic RBC partitioning occurring in the network. In total, 22 divergent bifurcations and 54 independent vessel segments (excluding 12 repetitive branches) are identified from the three ROIs. To quantify the distribution of RBCs at each divergent bifurcation (Fig. S2d–f), we introduce two variables following the practice of (47, 30): 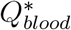, denoting the proportion of blood flow that a given child branch receives from its parent branch 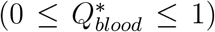; 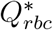, denoting the proportion of RBC flux likewise 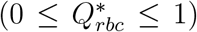. If 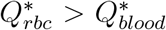, the child branch is receiving more RBCs than linear allocation and we define it as an “RBC-enriched” vessel; if 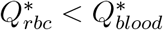, the child branch is receiving fewer RBCs than the linear hypothesis and we define it as an “RBC-depleted” vessel.

#### 5.3.2 Evaluation of simulation data against the phase separation model

Several models have been established to describe the phenomenon of plasma skimming (47, 75, 76, 77). Among others, the most widely-applied one is probably the empirical phase-separation model (hereafter referred to as PSM) developed by Pries and co-workers based on *in vivo* experiments and theoretical modelling (47, 48, 49, 78, 79). In brevity, the PSM derived a set of empirical equations from experimental observation of arteriolar bifurcations in rat mesentery, and established a flow-mediated mechanism to quantitatively describe the RBC fluxes received by child branches of diverging bifurcations within a microvascular network. The PSM correlates the fractional RBC flux *FQ_E_* in a child vessel of a divergent bifurcation with the fractional blood flow *FQ_B_* that it receives:

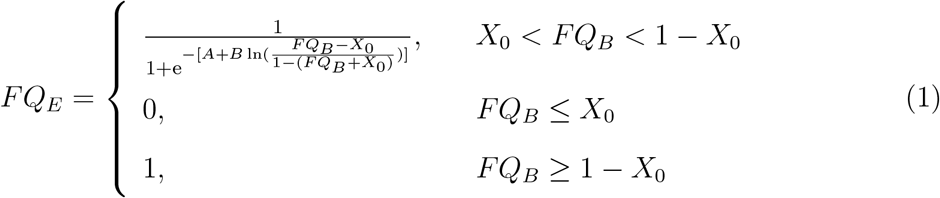

where *A*, *B* and *X*_0_ are fitting parameters derived *via* linear regression analysis. Physically, *A* reflects the size difference of the two child vessels, *B* reflects the shape of the haematocrit profile in the parent vessel and *X*_0_ is related to thickness of the cell-free layer near the corresponding vessel wall. We follow the original formulation in (49, 78):

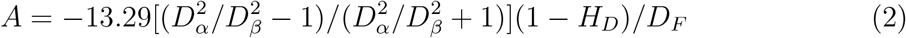

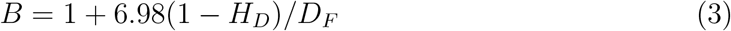

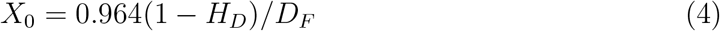

wherein *D_α_*, *D_β_* and *D_F_* are diameters of the two child branches and the parent branch (evaluated in *μ*m), respectively. *H_D_* is the discharge haematocrit of the parent branch.

With *D_α_*, *D_β_* and *D_F_* measured from the vascular geometry and *H_D_* obtained from the simulation (*via* an automated RBC-counting procedure), *A, B, X*_0_ can be calculated individually for each bifurcation and the empirical curves are plotted as *FQ_E_* versus *FQ_B_*. Equivalently, the simulation data are plotted in the form of 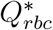 versus 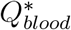 for comparison with the empirical predictions. The results for bifurcations in ROI-1, ROI-2 and ROI-3 are listed in Sec. S2 of the Supplementary Materials (Fig.s S3–S4 and Table S4). To evaluate the potential effect of cell volume difference as proposed in (80) between our RBC model (100 fl in volume) and realistic mouse RBCs (about 56.51 fl in volume), empirical predictions with rescaled fitting parameters *A*′, *B*′, *X*_0_′ are also examined:

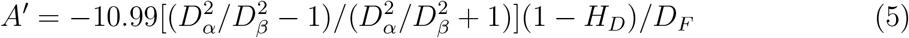

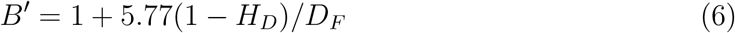

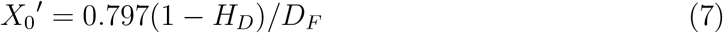

Quantitatively similar results to predictions by the original formulation (*i.e.* adopting *A, B, X*_0_) are obtained for all bifurcations in terms of 5% error threshold. The complete error evaluation against simulation data is in Table S5 of the Supplementary Materials.

#### 5.3.3 Statistical analysis

Statistical tests are conducted without any removal or modification of the original data. Testing of data normality is performed with the Shapiro-Wilk test and quantile-quantile plots. For normally or approximately normally distributed data, the parametric Welch’s T test (without assuming equal variances between two independent samples) is used and the means of data are measured, *e.g.* Figure 3i,j. For asymmetrically distributed data, the nonparametric Mann-Whitney U test is used and the medians of data are measured, *e.g.* Figure 1d,f and Figure 2d,e,f. All statistical tests are non-paired and two-sided. The difference between groups is considered statistically significant for \**P* < 0.05, very significant for \*\**P* < 0.01, and most significant for \*\*\**P* < 0.001; *P* > 0.05 is considered non-significant (ns).

## Supporting information

Supplementary Movie1_ROI-1

Supplementary Movie2_ROI-2

Supplementary Movie3_ROI-3

## 6 Acknowledgements

The authors would like to thank Aby Joseph and Jesse Schallek at University of Rochester for providing the *in vivo* data of RBC velocities/fluxes in mouse retina. We also acknowledge the contribution of the *HemeLB* development team. Q.Z. thanks the University of Edinburgh for the award of a Principal’s Career Development Scholarship and an Edinburgh Global Research Scholarship. T.P., I.F., L.T.E., H.G. and M.O.B. graciously acknowledge their funding as part of a Fondation Leducq Transatlantic Network of Excellence (17 CVD 03). T.K.’s and M.O.B.’s contributions have been funded through two Chancellor’s Fellowships at the University of Edinburgh. M.O.B. is supported by grants from EPSRC (EP/R029598/1, EP/R021600/1, EP/T008806/1), Fondation Leducq (17 CVD 03), and the European Union’s Horizon 2020 research and innovation programme under grant agreement No 801423. Supercomputing time on the ARCHER UK National Supercomputing Service (http://www.archer.ac.uk) was provided by the “UK Consortium on Mesoscale Engineering Sciences (UKCOMES)” under the EPSRC Grant No. EP/R029598/1. The authors declare that they have no competing financial interests.

## Supplementary Materials

### S1 Simulations of cellular blood flow in microvascular networks of mouse retina

**Fig. S1.**
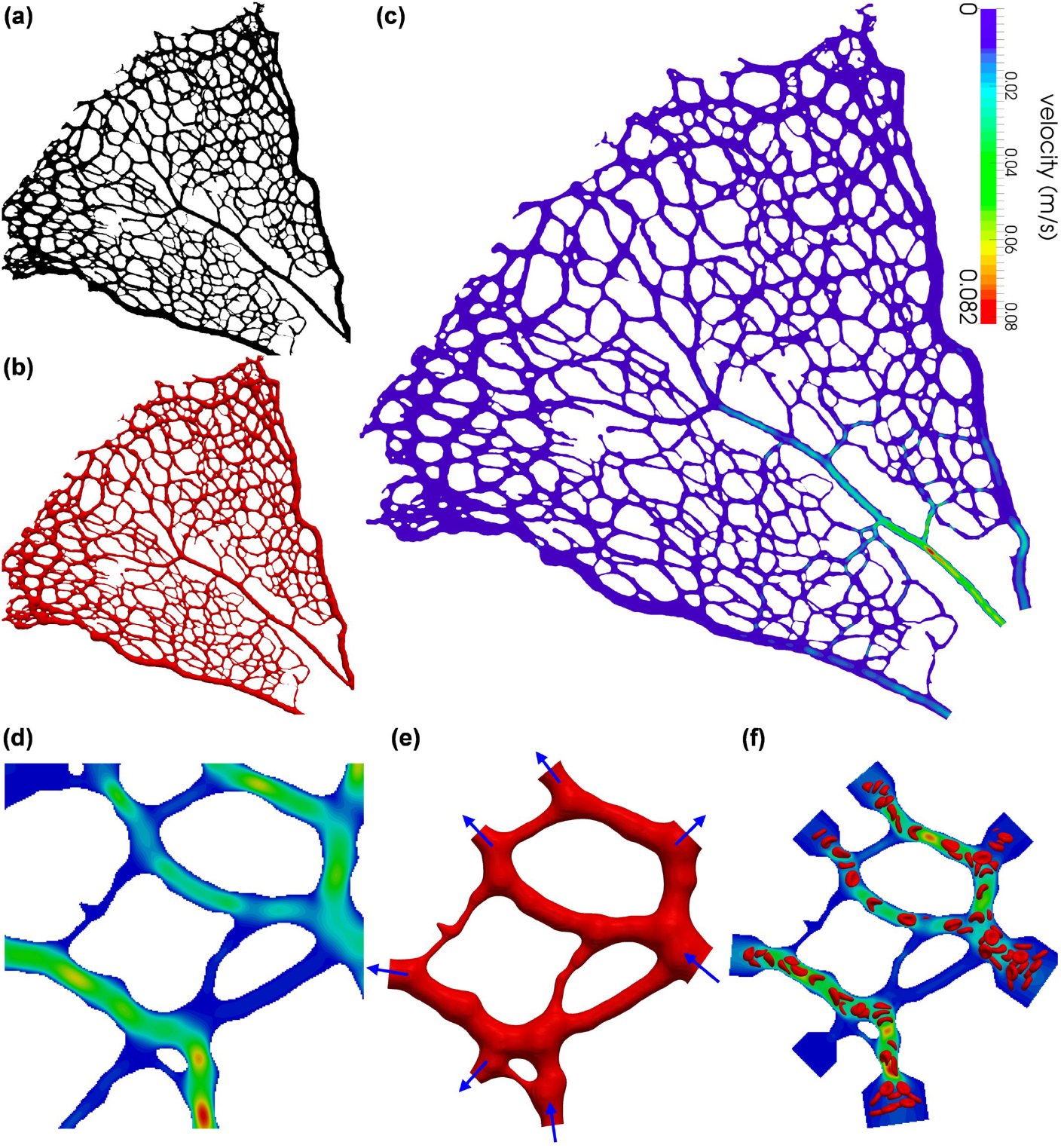
Procedure of model configuration and simulation setup for simulating cellular blood flow in designated regions of interest (ROIs) from the vascular plexus of a P5 mouse retina. (**a**) Binary image of the vascular plexus stained for Col.IV. (**b**) Reconstructed whole-network geometry of the vessel lumina surface from (a). (**c**) Velocity field within the retinal network resolved by a flow model applying the non-Newtonian Carreau-Yasuda (NNCY) blood rheology in (b). (**d**) Zoomed in velocity field for a designated ROI in (c). (**e**) Clipped ROI subset from the whole-network geometry with inlets/outlets indicated by the inward/outward arrows. (**f**) Cellular flow simulation in the designated ROI imposing inflow/outflow and pressure boundary conditions extracted from (d).

**Table S1.**
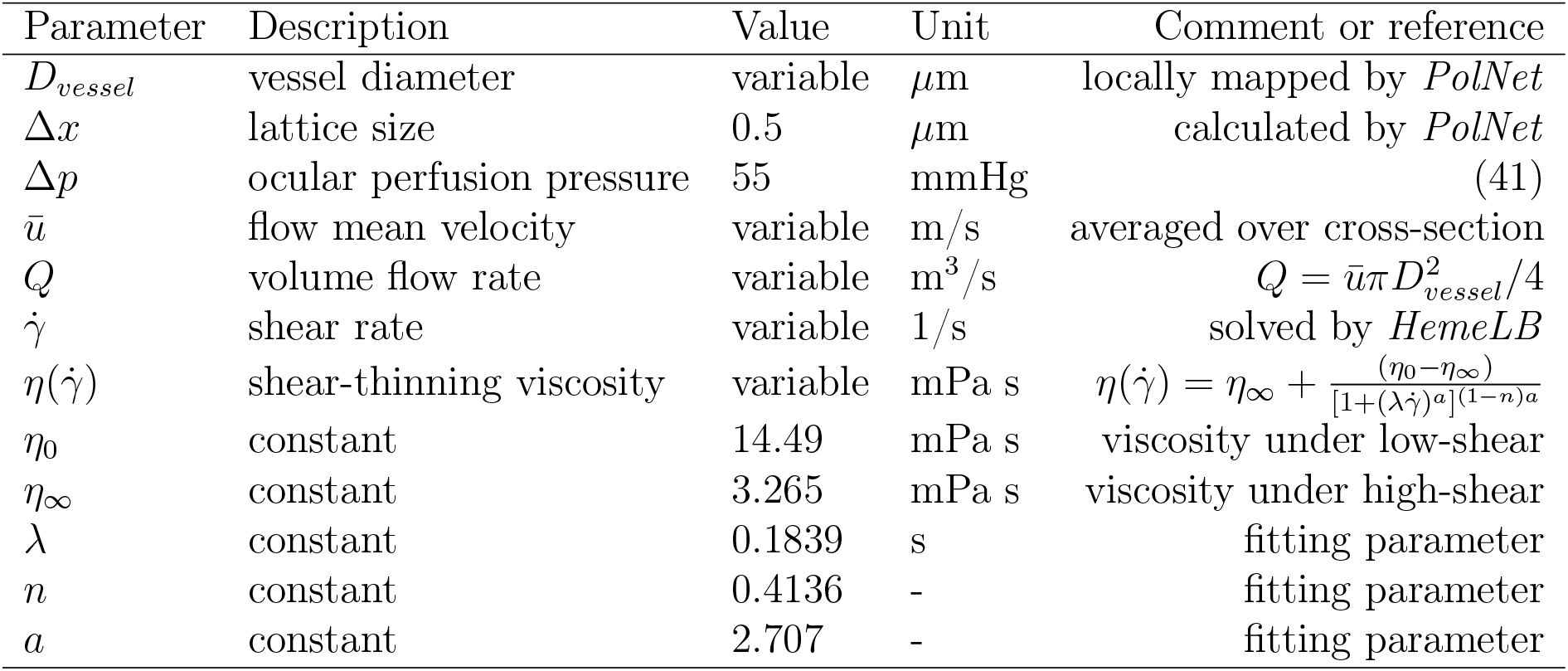
Key parameters of the whole-plexus simulation. A simplified rheology model, namely the non-Newtonian Carreau-Yasuda (NNCY) model (74, 41) is used.

The immersed-boundary-lattice-Boltzmann method (IB-LBM, (81)) is employed to model blood flow as a suspension of deformable RBCs. The fluid flow governed by the Navier-Stokes equations is solved by the lattice-Boltzmann method (LBM) with standard D3Q19 lattice (82), BGK collision operator (83) and Guo’s forcing scheme (84). The no-slip condition on vessel walls is implemented with the Bouzidi-Firdaouss-Lallemand method (BFL, 85). Open boundaries are handled with the Ladd implementation of velocity boundary conditions (86) for control of volume flow rates at multiple inlets and outlets. The RBCs are modelled as Lagrangian membranes using a finite element method (FEM). The fluid structure interaction (FSI) between the flow and the RBC membranes is realised using the immersed-boundary method (IBM, 87), which tackles the velocity interpolation and force spreading. The algorithm coupling the LBM, FEM and IBM is implemented in the open-source blood flow simulation software *HemeLB* (41; https://github.com/hemelb-codes/hemelb) for parallel computing.

Each RBC is modelled as a closed membrane consisting of *N_f_* triangular facets and present a discocyte shape at rest (81). The mesh resolution of the membrane (namely *N_f_*) matches the lattice size Δ*x* of the flow domain for numerical stability and accuracy (see 88 for a detailed numerical analysis). The RBC membrane is hyperelastic, isotropic and homogeneous. Its mechanical properties are controlled by several moduli (*κ*_*s*_, *κ*_*b*_, *κ*_*α*_, *κ*_*A*_, *κ*_*V*_) governing energy contributions from strain, bending, area and volume of the membrane (see Table S2). Enclosed by the RBC membrane is the cytosol treated as a Newtonian fluid of plasma viscosity. The viscosity of the RBC membrane itself is not considered in the present material model.

**Table S2.** Key parameters of the RBC model. The symbol “~” represents dimensionless simulation units. Please refer to (81) for full details of the RBC model.

| Parameter | Description | Value | Unit | Comment or reference |
| --- | --- | --- | --- | --- |
| $r_{rbc}$ | RBC radius | 4 | $\mu\text{m}$ | (89) |
| $A_{rbc}$ | RBC surface | 140 | $\mu\text{m}^2$ | (89) |
| $V_{rbc}$ | RBC volume | 100 | $\mu\text{m}^3$ | (89) |
| $\eta_{plasma}$ | plasma viscosity | 1 | mPa s | modelled as liquid water |
| $\eta_{cytosol}$ | artificial cytosol viscosity | 1 | mPa s | assuming $\eta_{cytosol} = \eta_{plasma}$ |
| $\Gamma_{nominal}$ | Föppl-von Kármán number | 400 | – | $\Gamma_{nominal} = \frac{\kappa_s r_{rbc}^2}{\kappa_b}$ |
| $N_f$ | number of facets | 1280 | – | $N_f = 20(\frac{r_{rbc}}{\Delta x})^2$ |
| $\kappa_s$ | strain modulus | 50 | $\mu\text{N/m}$ | resisting in-plane shearing |
| $\kappa_b$ | bending modulus | $2 \times 10^{-18}$ | Nm | resisting off-plane bending |
| $\tilde{\kappa}_\alpha$ | dilation modulus | 0.5 | – | conserving local area |
| $\tilde{\kappa}_A$ | surface modulus | 1 | – | conserving RBC surface |
| $\tilde{\kappa}_V$ | volume modulus | 1 | – | conserving RBC volume |

The morphological deformation of an RBC in small vessels or channels are known to be dominated by the strain modulus *κ_s_* and the bending modulus *κ_b_*, both of which have been extensively measured for healthy human RBCs using diverse experimental techniques. The commonly accepted values from different experiments are *κ_s_* = 5.5 ± 3.3 *μ*N/m and *κ_b_* = 1.15 ± 0.9 × 10^−19^ Nm, respectively (see reviews (89) and (90)). However, because these measurements all rely on certain deformation protocols that do not necessarily reflect the complex microcirculatory conditions, the obtained values may not apply to RBCs travelling in capillary networks of the mouse retina. Indeed, we find that the *κ_s_* and *κ_b_* required to maintain the integrity of the RBC membrane in our cellular simulations are roughly one order larger in magnitude than the reported values (see the simulation values in Table S2). This substantial increase in required *κ_s_* and *κ_b_* for reasonable RBC morphology may also arise from the intrinsic difference in the haemodynamic environment (*e.g.* magnitude ranges of the shear rate and wall shear stress) of the microcirculation system between mice and humans.

**Table S3.** Boundary conditions for RBC simulations in designated subsets of the retinal network, namely ROI-1, ROI-2 and ROI-3. *û* represents the centre-line velocity at the selected cross-section (circular) of a given vessel that serves as the inlet/outlet of the ROI and *p* is a reference pressure. *û* is set as positive for inlets and negative for outlets.

| ROI | Inlet/Outlet | Boundary type | Value | Unit | $D_{vessel} (\mu\text{m})$ |
| --- | --- | --- | --- | --- | --- |
| 1 | $\hat{u}_{in1}$ | velocity | 11.710 | mm/s | 8.0 |
| | $\hat{u}_{in2}$ | velocity | 3.756 | mm/s | 15.7 |
| | $\hat{u}_{out1}$ | velocity | -0.774 | mm/s | 5.4 |
| | $\hat{u}_{out2}$ | velocity | -6.164 | mm/s | 10.6 |
| | $\hat{u}_{out3}$ | velocity | -5.209 | mm/s | 7.1 |
| | $\hat{u}_{out4}$ | velocity | -4.829 | mm/s | 10.4 |
| | $p_{out5}$ | pressure | 0 | mmHg | 14.5 |
| 2 | $\hat{u}_{in1}$ | velocity | 4.222 | mm/s | 19.0 |
| | $\hat{u}_{in2}$ | velocity | 1.201 | mm/s | 5.5 |
| | $\hat{u}_{in3}$ | velocity | 6.367 | mm/s | 7.8 |
| | $\hat{u}_{out1}$ | velocity | -11.658 | mm/s | 10.3 |
| | $\hat{u}_{out2}$ | velocity | -7.182 | mm/s | 8.6 |
| | $\hat{u}_{out3}$ | velocity | -0.514 | mm/s | 7.3 |
| | $\hat{u}_{out4}$ | velocity | -0.113 | mm/s | 6.8 |
| | $p_{out5}$ | pressure | 0 | mmHg | 8.9 |
| 3 | $\hat{u}_{in1}$ | velocity | 1.171 | mm/s | 7.1 |
| | $\hat{u}_{in2}$ | velocity | 10.277 | mm/s | 13.5 |
| | $\hat{u}_{in3}$ | velocity | 0.049 | mm/s | 6.9 |
| | $\hat{u}_{in4}$ | velocity | 0.002 | mm/s | 2.4 |
| | $\hat{u}_{out1}$ | velocity | -14.913 | mm/s | 8.8 |
| | $\hat{u}_{out2}$ | velocity | -30.298 | mm/s | 4.7 |
| | $p_{out3}$ | pressure | 0 | mmHg | 2.2 |

### S2 Evaluation of simulation data against empirical model

**Fig. S2.**
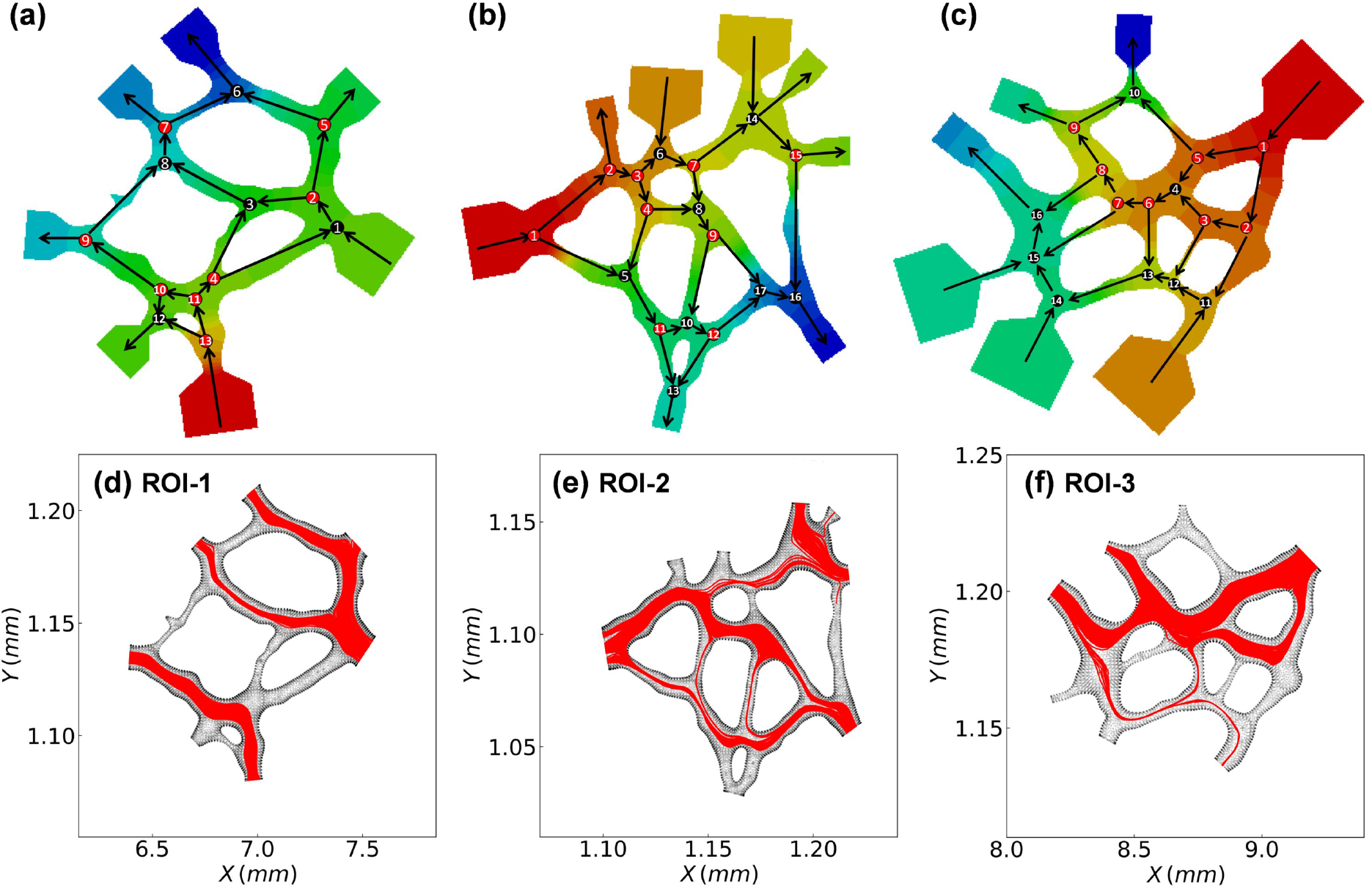
Quantification of RBC flow in designated subsets of the retinal network. (**a**–**c**) Flow patterns detected within ROI-1, ROI-2 and ROI-3, respectively. The arrows within the ROIs indicate the flow directions in individual vessel segments, and the background contour indicates the pressure field, with a warmer colour (*e.g.* red) representing a higher pressure. The divergent bifurcations within each ROI are marked with red circles and the convergent ones with black. (**d**–**f**) Combined cell trajectories in the ROIs throughout RBC simulations lasting for 0.33 s each.

**Fig. S3.**
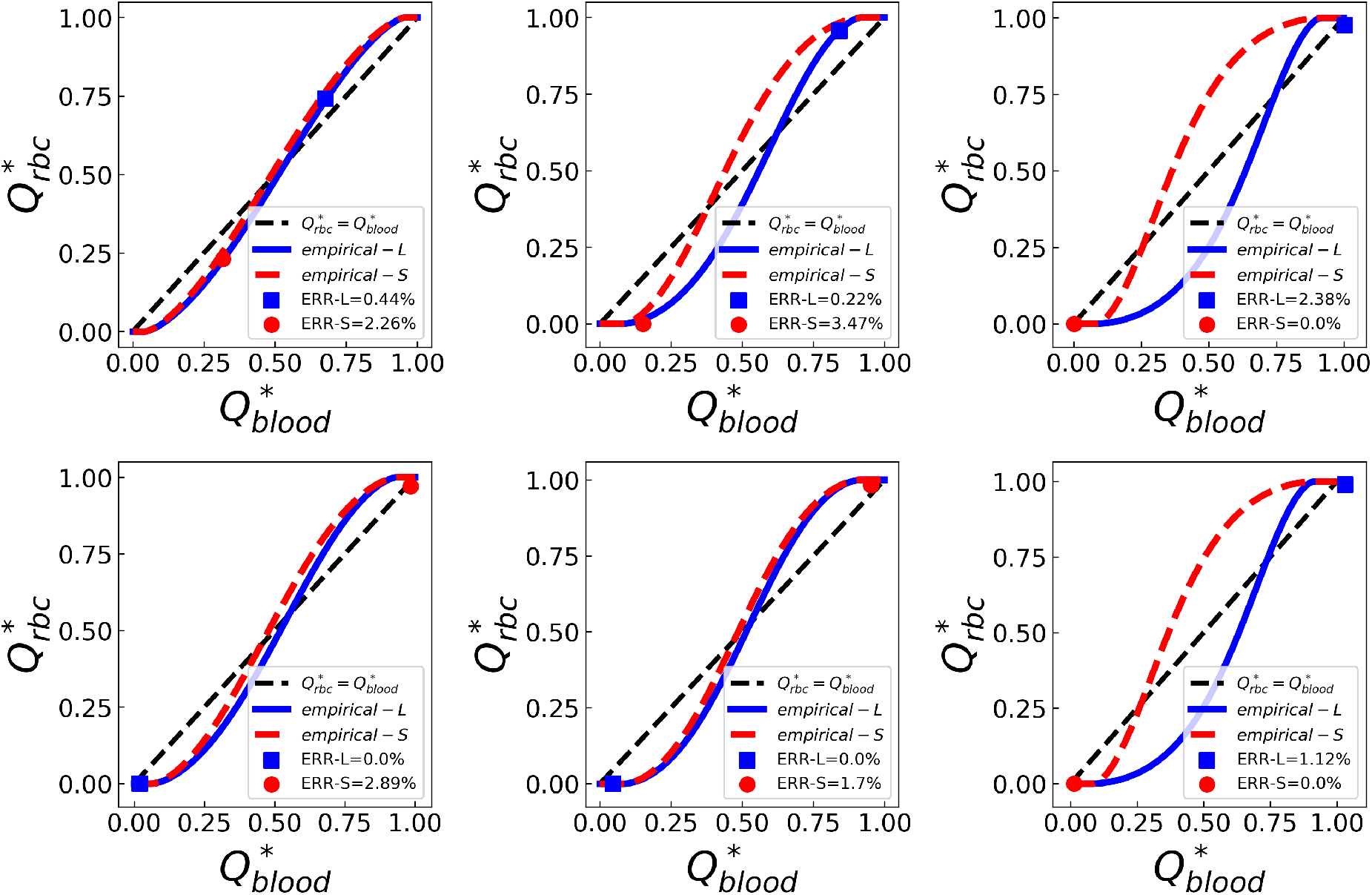
Evaluation of simulation data against empirical predictions by the phase separation model (49). Simulation data extracted from divergent bifurcations in ROI-1 (see Fig. S2d) are plotted as fractional RBC flux 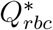 against fractional blood flow 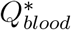. The simulation data are represented by squares/circles and the empirical predictions by solid lines. In each bifurcation, the relatively larger child branch is termed “L” and relatively smaller child branch termed “S”. The black dashed line represent a linear hypothesis for 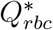 and 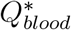 in the absence of plasma skimming.

**Table S4.**
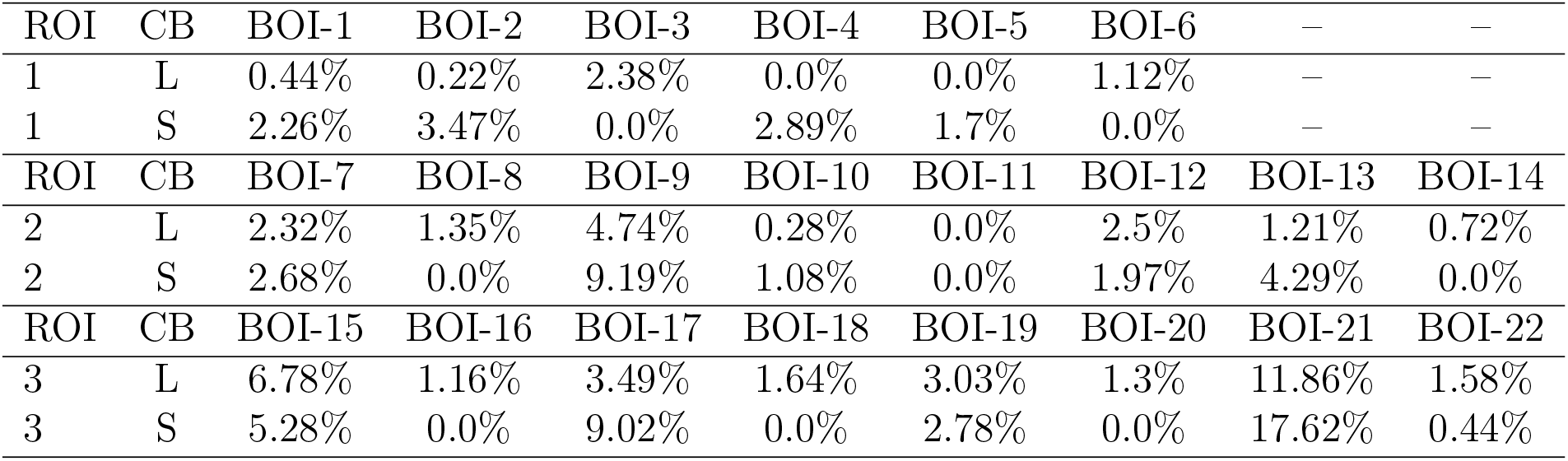
Errors of empirical prediction by the phase separation model (49) in comparison to our simulation data. “ROI” represents region of interest; “BOI” represents bifurcation of interest; “CB” represents child branch. “L” and “S” refer to the relatively larger and smaller child branch within each bifurcation, respectively.

| ROI | CB | BOI-1 | BOI-2 | BOI-3 | BOI-4 | BOI-5 | BOI-6 | — | — |
| --- | --- | --- | --- | --- | --- | --- | --- | --- | --- |
| 1 | L | 0.44% | 0.22% | 2.38% | 0.0% | 0.0% | 1.12% | — | — |
| 1 | S | 2.26% | 3.47% | 0.0% | 2.89% | 1.7% | 0.0% | — | — |
| ROI | CB | BOI-7 | BOI-8 | BOI-9 | BOI-10 | BOI-11 | BOI-12 | BOI-13 | BOI-14 |
| 2 | L | 2.32% | 1.35% | 4.74% | 0.28% | 0.0% | 2.5% | 1.21% | 0.72% |
| 2 | S | 2.68% | 0.0% | 9.19% | 1.08% | 0.0% | 1.97% | 4.29% | 0.0% |
| ROI | CB | BOI-15 | BOI-16 | BOI-17 | BOI-18 | BOI-19 | BOI-20 | BOI-21 | BOI-22 |
| 3 | L | 6.78% | 1.16% | 3.49% | 1.64% | 3.03% | 1.3% | 11.86% | 1.58% |
| 3 | S | 5.28% | 0.0% | 9.02% | 0.0% | 2.78% | 0.0% | 17.62% | 0.44% |

**Fig. S4.**
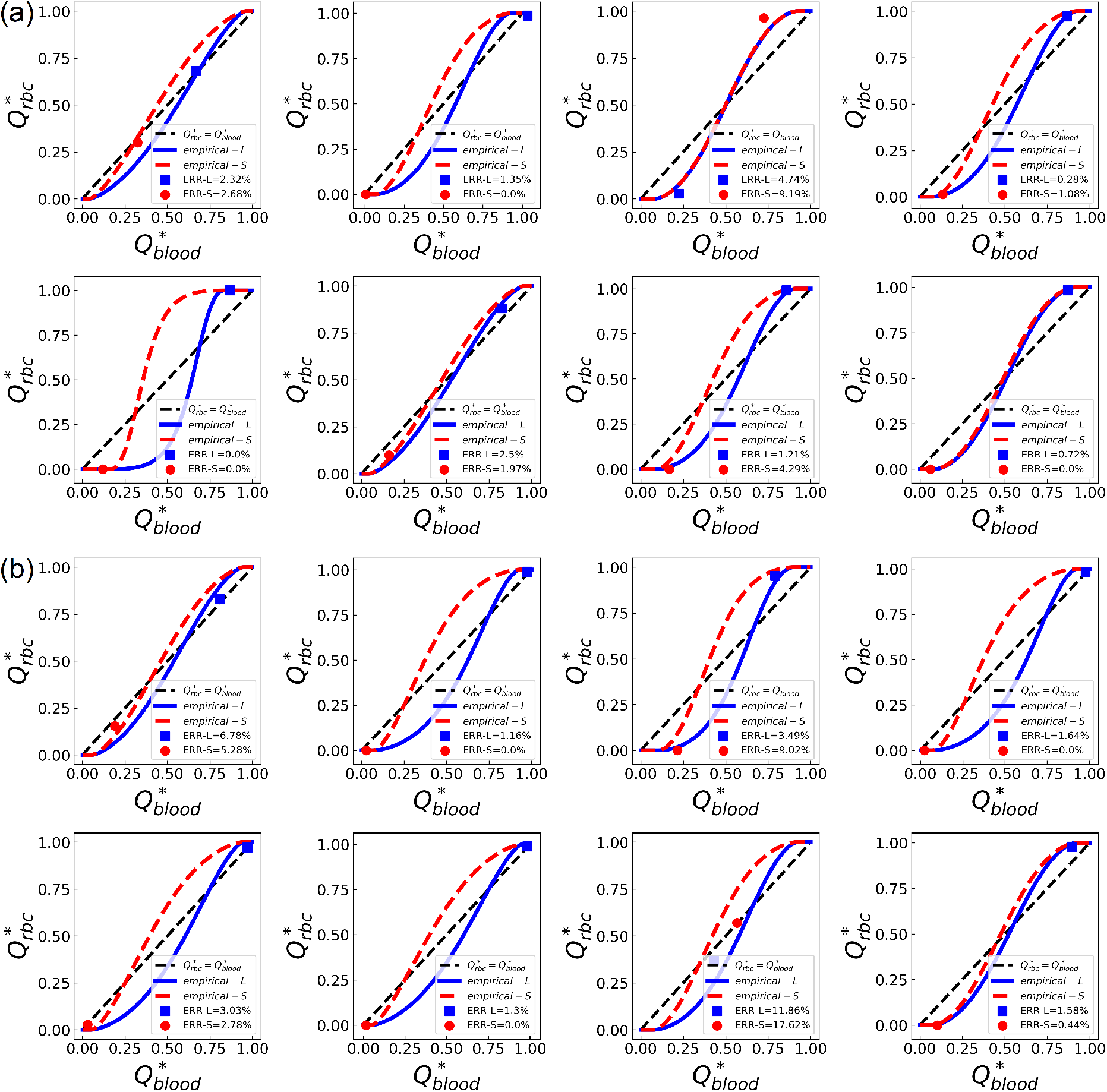
Evaluation of simulation data against empirical predictions. (a) Same caption as Fig. S3, except for ROI-2 (see Fig. S2e).(b) Same caption as Fig. S3, except for ROI-3 (see Fig. S2f).

**Table S5.** Same caption as Table S4, except that the phase separation model here adopts revised parameters *A*′, *B*′, *X*_0_′ based on (49). *A*′, *B*′, *X*_0_′ account for the RBC volume difference (see Equations (5)–(7)).

| ROI | CB | BOI-1 | BOI-2 | BOI-3 | BOI-4 | BOI-5 | BOI-6 | – | – |
| --- | --- | --- | --- | --- | --- | --- | --- | --- | --- |
| 1 | L | 0.93% | 1.94% | 2.38% | 0.0% | 0.0% | 1.12% | – | – |
| 1 | S | 0.85% | 1.89% | 0.0% | 2.89% | 1.7% | 0.0% | – | – |
| ROI | CB | BOI-7 | BOI-8 | BOI-9 | BOI-10 | BOI-11 | BOI-12 | BOI-13 | BOI-14 |
| 2 | L | 2.3% | 1.35% | 2.55% | 1.64% | 0.0% | 4.03% | 0.15% | 1.44% |
| 2 | S | 2.54% | 0.0% | 6.54% | 0.26% | 0.0% | 3.44% | 2.35% | 0.0% |
| ROI | CB | BOI-15 | BOI-16 | BOI-17 | BOI-18 | BOI-19 | BOI-20 | BOI-21 | BOI-22 |
| 3 | L | 8.54% | 1.16% | 0.49% | 1.64% | 3.03% | 1.3% | 15.15% | 2.07% |
| 3 | S | 7.04% | 0.0% | 5.94% | 0.0% | 2.78% | 0.0% | 20.9% | 0.04% |

### S3 Asymmetry of velocity profile in RBC flow

**Fig. S5.**
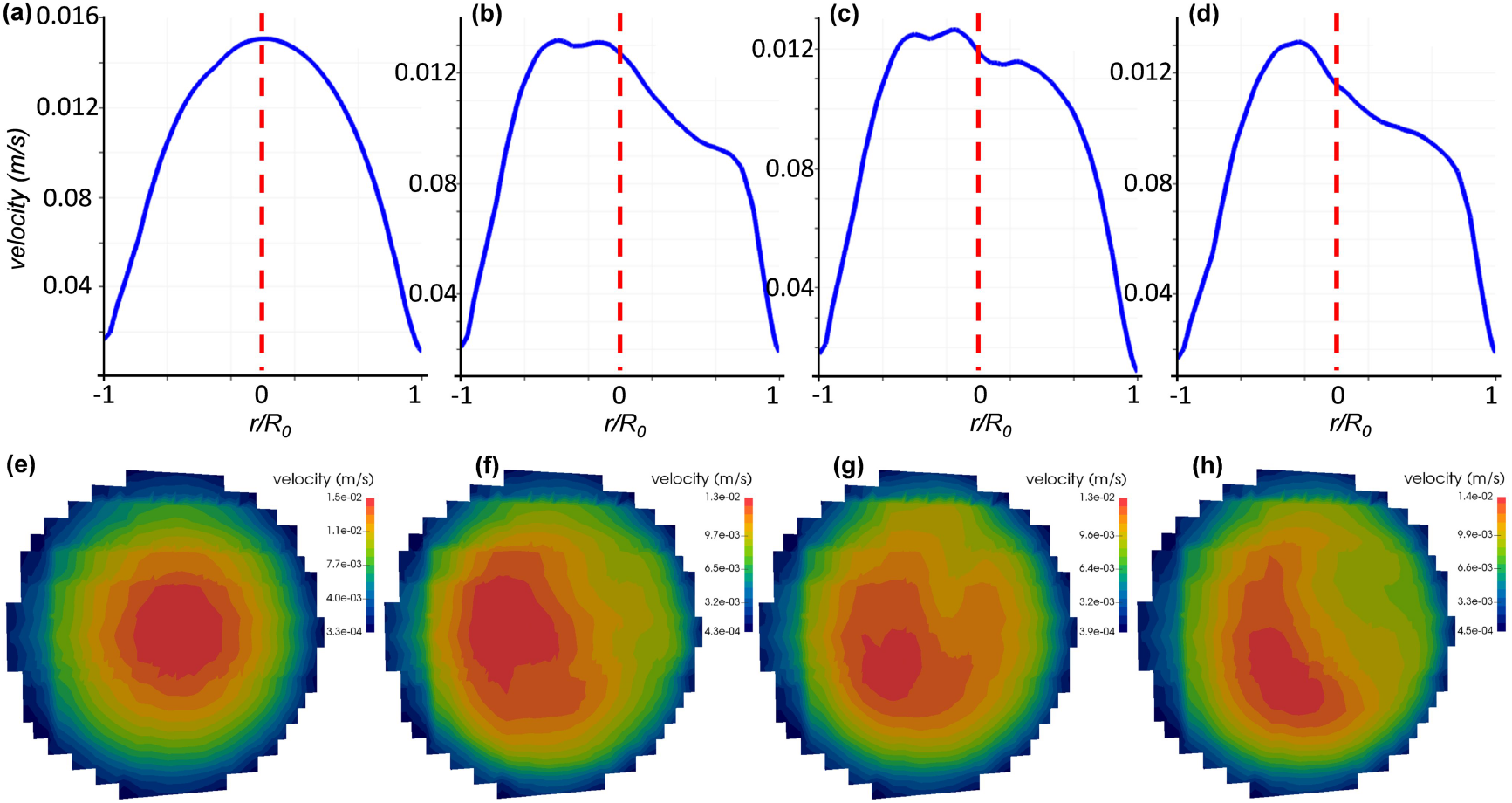
Temporal velocity profile and flow pattern in the RBC flow. (**a**-**d**) Velocity profiles over time at the red solid line labelled in Figure 6d. (**e**-**h**) Corresponding cross-sectional velocity contours at the same position. (a) and (e): *t*_1_ = 0.042s. (b) and (f): *t*_2_ = 0.125s. (c) and (g): *t*_3_ = 0.208s. (d) and (h): *t*_4_ = 0.291s.

**Movie S1.** Video of the simulated RBC flow in region of interest ROI-1 (see Fig. 1).

**Movie S2.** Video of the simulated RBC flow in region of interest ROI-2 (see Fig. 1).

**Movie S3.** Video of the simulated RBC flow in region of interest ROI-3 (see Fig. 1).

